# Leveraging big data for classification of children who stutter from fluent peers

**DOI:** 10.1101/2020.10.28.359711

**Authors:** Saige Rutherford, Mike Angstadt, Chandra Sripada, Soo-Eun Chang

**Author notes:** **Corresponding author:** Saige Rutherford, Address: 4250 Plymouth Rd, Ann Arbor, MI 48109.

## Abstract

**Introduction:** Large datasets, consisting of hundreds or thousands of subjects, are becoming the new data standard within the neuroimaging community. While big data creates numerous benefits, such as detecting smaller effects, many of these big datasets have focused on non-clinical populations. The heterogeneity of clinical populations makes creating datasets of equal size and quality more challenging. There is a need for methods to connect these robust large datasets with the carefully curated clinical datasets collected over the past decades.

**Methods:** In this study, resting-state fMRI data from the Adolescent Brain Cognitive Development study (N=1509) and the Human Connectome Project (N=910) is used to discover generalizable brain features for use in an out-of-sample (N=121) multivariate predictive model to classify young (3-10yrs) children who stutter from fluent peers.

**Results:** Accuracy up to 72% classification is achieved using 10-fold cross validation. This study suggests that big data has the potential to yield generalizable biomarkers that are clinically meaningful. Specifically, this is the first study to demonstrate that big data-derived brain features can differentiate children who stutter from their fluent peers and provide novel information on brain networks relevant to stuttering pathophysiology.

**Discussion:** The results provide a significant expansion to previous understanding of the neural bases of stuttering. In addition to auditory, somatomotor, and subcortical networks, the big data-based models highlight the importance of considering large scale brain networks supporting error sensitivity, attention, cognitive control, and emotion regulation/self-inspection in the neural bases of stuttering.

## Introduction

Childhood onset fluency disorder, more commonly known as stuttering, is a neurodevelopmental disorder that affects 1% of the general population (5-8% of all preschool-age children) (Yairi & Ambrose, 2013). Stuttering significantly impedes the speaker’s ability to produce the rhythmic, fluid flow of speech sounds, and can lead to substantial negative psychosocial consequences.

Most studies examining stuttering have applied methodological approaches that focus on specific hypothesis driven brain regions or examine connectivity among a few areas of interest (See for reviews (Chang et al., 2019; Neef et al., 2015)) However, as a complex neurodevelopmental disorder, stuttering likely emerges with subtle changes in large-scale network connections that support multiple functions, including cognitive/language, attention, and motor control. In a recent study, we used a connectomics approach to examine intra- and inter-network connectivity of large-scale intrinsic connectivity networks for the first time to examine stuttering children. We found that children who stutter, regardless of later persistence or recovery from stuttering, could be differentiated from their non-stuttering peers based on earlier collected resting-state fMRI scans (Chang et al., 2018). In general, somatomotor network connectivity was aberrant in children who stutter. However, the differences were also reflected in the somatomotor network’s connectivity with other large-scale networks such as attention and default mode networks. This study also reported network connectivity patterns that differentiated persistently stuttering children from recovered children. Given the lack of a held-out test set or cross validated model performance, these results warrant replication and expansion through larger out of sample predictive studies.

A limitation of most clinical studies, especially those involving scanning young children, is that the sample sizes tend to be small, leading to limited statistical power to discover true effects and prone to finding false positives and lack of replication. Due to the small sample sizes, prediction-based (as opposed to associative) studies are rare. Reflecting on these issues’ seriousness, big datasets consisting of hundreds or thousands of subjects are becoming the new data standard within the neuroimaging community. Big data creates numerous benefits, including allowing for more ambitious statistical analyses than smaller studies due to the increased power and better estimations of model generalizability. One important benefit of big datasets is that underlying “features” inherent in the dataset, such as in specific connectivity patterns of large-scale networks extracted from resting-state fMRI data, may be more reliably measured, and modes of variation of these features are better estimated due to the increased power.

Previous studies have shown that certain brain features can be linked to phenotypes of interest (e.g., variations in IQ, attention, etc.), and can also be used for the prediction of cognitive or clinical phenotypes (Beaty et al., 2018; Chen et al., 2020; Dubois et al., 2018; Finn et al., 2015; Goñi et al., 2014; He et al., 2020; Kong et al., 2019; Lake et al., 2019; Nguyen et al., 2020; Rosenberg et al., 2015, 2020; Sripada, Angstadt, Rutherford, Taxali, Clark, et al., 2020; Weis et al., 2020; Wu et al., 2020). Recent work in the ABCD study showed that until sample sizes approach thousands of subjects, brain-behavior relationships were underpowered and statistical errors were inflated (Marek et al., 2020). The rich research questions regarding whether big data-derived brain features can be applied to smaller clinical datasets have not yet been explored. Other work has suggested that big-data applied to small-data, dubbed “meta-matching,” is potentially useful but did not explore across dataset predictive model transfer (He et al., 2020). There is a strong need for methods that connect these powerful large datasets with the carefully curated clinical datasets. We address the feasibility of across dataset model transfer by discovering brain features in the ABCD and HCP studies and applying them to a smaller (out of sample) clinical dataset.

This study aimed to create an analysis framework to combine big data with a more modest sample size clinical dataset. We leveraged a multivariate predictive modeling method, brain basis set (Sripada, Angstadt, et al., 2019; Sripada, Angstadt, Rutherford, Taxali, & Shedden, 2020; Sripada, Rutherford, et al., 2019), to bring the power of big data into a framework examining group differences in brain connectivity present in children who stutter. Our pipeline begins with feature discovery and selection in large open datasets, using resting-state functional MRI data from the Human Connectome Project (HCP) and the Adolescent Brain Cognitive Development (ABCD) study. We then transfer these big-data brain features to an out-of-sample clinical dataset, consisting of resting-state fMRI data collected from children who stutter and healthy controls. These data were collected as part of an on-going longitudinal study in stuttering (Chang et al., 2018; Garnett et al., 2018).

This modeling approach fuses unsupervised and supervised learning techniques. The initial decomposition of fMRI data (feature discovery) is unsupervised through principal component analysis (PCA), meaning it is unaware of the data’s behavioral characteristics. The predictive modeling portion of the pipeline, which uses cross-validated logistic regression, is supervised because the model is informed (in the training set) of all participants’ clinical labels. Brain basis set takes advantage of the fact that, though functional connectomes are massive, complex objects, there is high redundancy in the set of connections that differ across people. This allows for a distilled set of PCA components to capture the most meaningful inter-individual variations and provide generalizability allowing us to use the basis set developed in one group for prediction in a separate clinical dataset.

In the big datasets used here for feature discovery, participants of both HCP and ABCD were tested on a battery of assessments, including those relevant to language and attention. Prior work using brain basis set modeling within these datasets showed the best predictive performance when predicting cognitive phenotypes, such as fluid intelligence or latent cognitive variables such as general cognitive ability (Sripada, Taxali, et al., 2019; Sui et al., 2020). Examining brain basis sets associated with behaviors relevant to children who stutter and applying them to an out of sample stuttering dataset may provide a way to predict subgroups within the stuttering group, such as categorizing those most likely to recover from stuttering or go on to develop chronic stuttering. Early prediction of the clinical population’s different clinical trajectories is important because it could prioritize clinical resources toward delivering early intervention to those children most vulnerable to developing persistent stuttering. Apart from clinical implications, better classification of children who stutter from their non-stuttering peers is likely to provide a breakthrough in understanding the complex neural bases of stuttering.

This work’s central goal is to test if big datasets, such as the ABCD study and the HCP data, can help discover a “better” brain basis set, which is a basis set that improves out of sample classification between children who stutter and fluent peers. The rationale behind this hypothesis is that larger sample sizes tend to discover more generalizable brain features. HCP and ABCD data contain higher quality MRI data (spatial and temporal resolution, longer scan length) than most clinical datasets (Casey et al., 2018; Van Essen et al., 2013). To test this big data prediction hypothesis, we directly compare the big data model’s performance to within-sample feature discovery-based models.

Predictive modeling work within the clinical neuroimaging community is often met with skepticism (Bzdok & Meyer-Lindenberg, 2018; Cabitza et al., 2017; Feczko et al., 2019; Lasko et al., 2017; Stephan, Bach, et al., 2016, p. 1; Stephan, Binder, et al., 2016). Much of the criticism surrounding predictive modeling stems from the fact that many predictive models are “black boxes,” yielding low interpretability in terms of the circuits involved (Rudin & Radin, 2019). We emphasize the importance of interpretable and *plausible* prediction in clinical samples, making these characteristics top priority in this work. Therefore, we included an additional analysis to move our work beyond broad statements about patients differing from healthy controls to characterizing which brain networks may contribute the most to the observed differences with the hope of informing future interventions.

## Materials & Methods

### Data acquisition

**<u>HCP</u>** All subjects and data were from the HCP-1200 release (Van Essen et al., 2013). Four runs of resting-state fMRI data (14.4 minutes each; two runs per day over two days) were acquired (TR = 720 ms).

**<u>ABCD</u>** Data from the curated ABCD annual release 1.1 were used, and full details are described in (Hagler et al., 2019). Imaging protocols were harmonized across sites and scanners. High spatial (2.4 mm isotropic) and temporal resolution (TR = 800 ms) resting-state fMRI was acquired in four separate runs (5min per run, 20 minutes total).

**<u>Stuttering</u>** During the rsfMRI scan, children lay supine with their eyes open. They were instructed to remain as still as possible. Preceding the MRI scanning session, all children were trained during a separate visit with a mock scanner to familiarize and desensitize them to the sights and sounds of the scanner and to practice being still inside the scanner bore (Chang et al., 2015, 2016). To ensure that the child remained calm and to minimize the possibility of movement, an experimenter sat by the child throughout the scan. MRI scans were acquired on a GE 3T Signa HDx scanner (GE Healthcare). Thirty-six contiguous 3-mm axial slices were collected with a gradient-echo EPI sequence (7 min) in an interleaved order (TR = 2500 ms). All procedures used in this study were approved by the Michigan State University Institutional Review Board. Informed consent was obtained according to the Declaration of Helsinki. All children were paid a nominal remuneration, and were given small prizes (e.g. stickers) for their participation. This study is not a clinical trial.

### In/Exclusion criteria

**<u>HCP</u>** subjects were eligible to be included if they had structural T1w and T2w data and had four complete resting-state fMRI runs (14m 24s each; 1206 subjects total in release files, 1003 with full resting state and structural). Subjects with more than 10% of frames censored were excluded from further analysis, and if there was incomplete phenotypic data, leaving 910 subjects.

**<u>ABCD</u>** subjects were eligible to be included if they had at least 4 minutes of good data (after motion censoring at FD>0.5mm) and a usable T1w image (*n*=2757). To remove unwanted sources of dependence in the dataset, only one sibling was randomly chosen to be retained for any family with more than one sibling (*n*=2494). Incomplete neurocognitive data were also criteria for exclusion, as were preprocessing errors in applying the field-maps. This left 1,509 subjects.

**<u>Stuttering</u>** subjects were eligible to be included if they had at least 4 minutes of good data (after motion censoring at FD>0.5mm) and a usable T1w image (*n*=121). The demographics of subjects included in our analysis are shown in Table 1.

**Table 1.** Participant demographics. Mean (standard deviation).

|  | **Stuttering** | **ABCD** | **HCP** |
| --- | --- | --- | --- |
| Sample size | 121 | 1509 | 910 |
| Age (years) | $6.14 \pm 1.9$ | $10.05 \pm 0.60$ | $28.5 \pm 3.7$ |
| Sex (M/F) | 61/60 | 783/726 | 435/475 |
| mean FD (mm) | $0.39 \pm 0.34$ | $0.21 \pm 0.08$ | $0.15 \pm 0.04$ |
| Percent stuttered syllables (%SLD) | $3.66 \pm 3.73$ | - | - |
| Stuttering severity instrument (SSI)<sup>1</sup> | $9.3 \pm 10.2$ | - | - |

### Data Preprocessing

We harmonized data preprocessing as much as possible across all three datasets used in this study. However, due to the nature of these datasets, small differences in data preprocessing occurred across these datasets, and each preprocessing workflow is described as follows.

**<u>HCP</u>** Processed volumetric data from the HCP minimal preprocessing pipeline, including ICA-FIX denoising, were used. Full details of these steps can be found in Glasser (Glasser et al., 2013) and Salimi-Korshidi (Salimi-Khorshidi et al., 2014). Briefly, T1w and T2w data were corrected for gradient-nonlinearity and readout distortions, inhomogeneity corrected and registered linearly and nonlinearly to MNI space using FSL’s FLIRT and FNIRT. BOLD fMRI data were also gradient-nonlinearity distortion corrected, rigidly realigned to adjust for motion, fieldmap corrected, aligned to the structural images, and then registered to MNI space with the nonlinear warping calculated from the structural images. Then FIX was applied to the data to identify and remove motion and other artifacts in the time-series. Images were smoothed with a 6mm Gaussian kernel and then resampled to 3mm isotropic resolution. The smoothed images then went through several resting-state processing steps, including a motion artifact removal steps comparable to the type B (i.e., recommended) stream of Siegel et al. (Siegel et al., 2017). These steps include linear detrending, CompCor to extract, and regress out the top 5 principal components of white matter and CSF (Behzadi et al., 2007), bandpass filtering from 0.1-0.01Hz, and motion scrubbing of frames that exceed a framewise displacement of 0.5mm.

**<u>ABCD</u>** Minimally preprocessed resting-state fMRI was used from data release 1.1. This data reflects the application of the following steps: i) gradient-nonlinearity distortions and inhomogeneity correction for structural data; and ii) gradient-nonlinearity distortion correction, rigid realignment to adjust for motion, and field map correction for functional data. Additional processing steps were applied by our group using SPM12, including co-registration using the CAT12 toolbox application, smoothing with a 6mm Gaussian kernel, and application of ICA-AROMA (Pruim et al., 2015). Resting-state processing steps were then applied, including linear detrending, CompCor (Behzadi et al., 2007), bandpass filtering from 0.1-0.01Hz, and motion scrubbing of frames that exceed a framewise displacement of 0.5mm.

**<u>Stuttering</u>** Data were processed using typical methods in Statistical Parametric Mapping (SPM12, Wellcome Institute of Cognitive Neurology, London). Slice time was corrected using sinc-interpolation, and all scans were realigned to the 10th volume acquired during each scan. Time-series of functional volumes were then co-registered with a high-resolution T1 image, spatially normalized to the MNI152 brain using the CAT12 toolbox, and then spatially smoothed with a 6 mm isotropic Gaussian kernel. ICA-AROMA (Pruim et al., 2015) was applied to the smoothed data for motion denoising. Resting-state processing steps were then applied, including linear detrending, CompCor (Behzadi et al., 2007), bandpass filtering from 0.1-0.01Hz, and motion scrubbing of frames that exceed a framewise displacement of 0.5mm.

### Connectome Generation

We calculated spatially averaged time series for each of 264 4.24mm radii ROIs from the parcellation of Power et al. (Power et al., 2011). We then calculated Pearson’s correlation coefficients between each ROI. These were then transformed using Fisher’s r to z-transformation. Connectomes are symmetric matrices that do not contain directionality information. Therefore, we vectorize each subject’s connectome’s upper triangle to create a 1 × 34,716 (264 choose 2) vector. All subject’s connectome vectors are then stacked, creating an *n* subjects × *p* connections matrix, where rows represent unique subjects and columns represent unique connections.

### Brain Basis Set Predictive Modeling

Brain Basis Set (BBS) is a multivariate predictive method that uses dimensionality reduction to produce a basis set of components to make phenotypic predictions (see Figure 1 for an overview). First, for the dimensionality reduction step, we submitted an *n* subjects *x p* connections matrix for both the HCP and ABCD training datasets (separately) for principal components analysis. Next, we moved to the stuttering dataset to calculate the expression scores for each of the k components for each subject by projecting each subject’s connectivity matrix onto each principal component. We then fit a logistic regression model with these expression scores as predictors and the phenotype of interest (the clinical diagnosis of stuttering) as the outcome. In a test dataset, we again calculated the expression scores for each component in the basis set for each test subject. We repeated this model within the stuttering dataset using 10-fold cross validation. Importantly, our model controls for nuisance variables (age, sex, linear and quadratic effects of motion).

**Figure 1.**
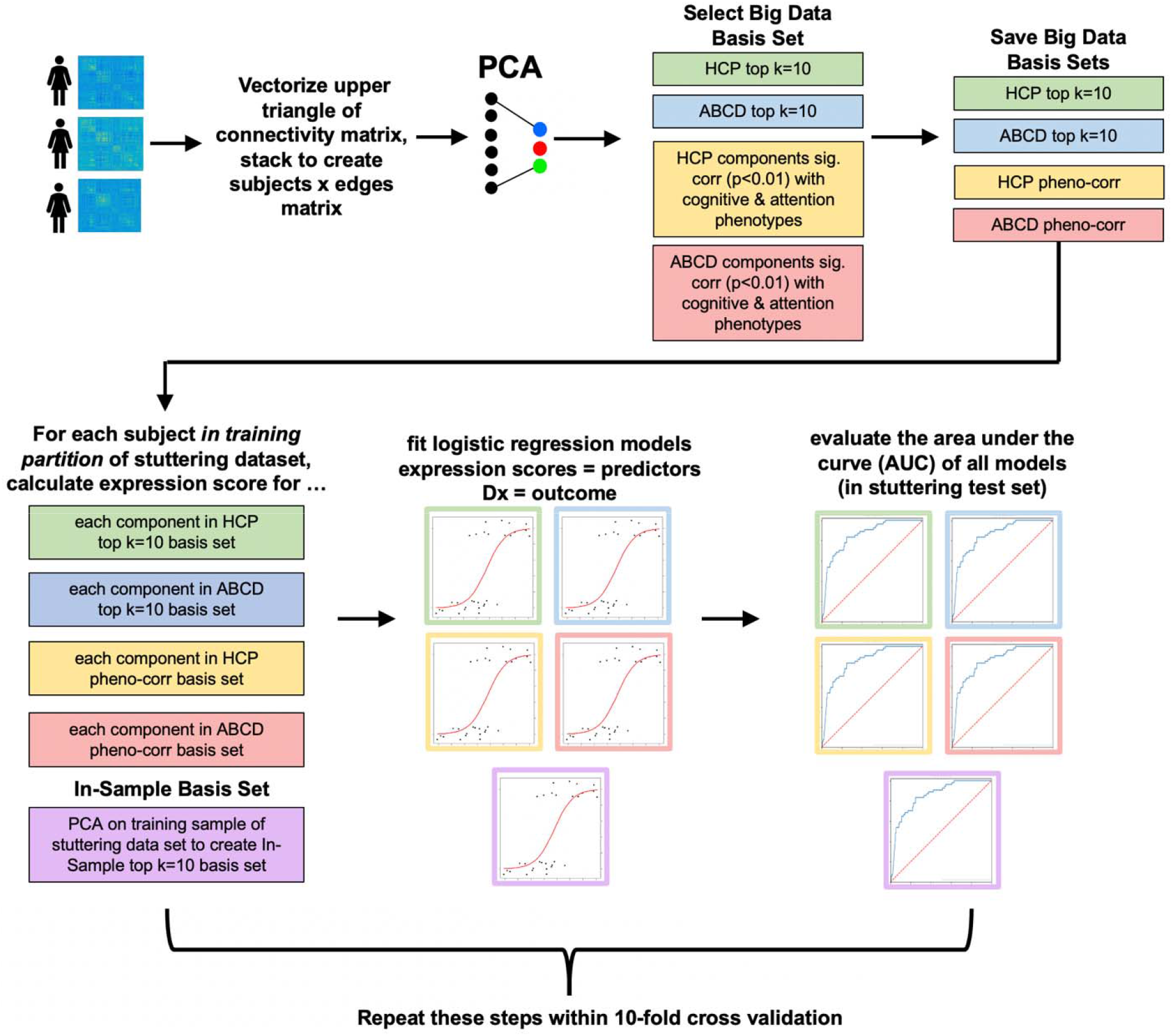
Overview of Brain Basis Set (BBS) Predictive Modeling. BBS utilizes dimensionality reduction with principal components analysis (PCA) to construct a high-quality feature set in a large data set (i.e., HCP & ABCD) and then apply the basis set to an out-of-sample clinical data set to the classification of children who stutter from fluent peers and compare this model performance to within-sample basis set using 10-fold cross validation.

### Feature Selection: Top 10 components (“Top k”) and phenotype-correlation (“Pheno-corr”) models

We tested three different techniques for selecting which components to use in the predictive model. First, we used the top k variance explaining components from the HCP basis set and then also from the ABCD basis set. Prior work using BBS modeling in ABCD and HCP datasets showed that somewhere between 50 to 100 components yields an optimal prediction of a broad array of behavioral phenotypes (Sripada, Angstadt, et al., 2019). However, due to the smaller sample size of the stuttering dataset (n=121) compared to the HCP (n=910) and ABCD (n=1509) sample sizes, we lowered this number due to the possibility of overfitting the data. The sample size of the clinical dataset is approximately 1/10^th^ the sample size of HCP and ABCD. Therefore, in this study, we used 1/10^th^ the number of components used in previous HCP & ABCD predictive models and set k equal to 10. These models are referred to as “HCP top k=10” and “ABCD top k=10”.

Next, HCP and ABCD basis set components that were significantly correlated (p<0.01) with the phenotypes of interest (cognitive variables from the NIHToolbox and sustained attention) were selected. These models are referred to as “ABCD pheno-corr” and “HCP pheno-corr” models. Cognitive NIHToolbox phenotypes were selected based on previous work in HCP and ABCD predictive modeling demonstrating that these cognitive phenotypes tend to yield the highest accuracy and test-retest reliability (Sripada, Taxali, et al., 2019). The choice of sustained attention among the phenotypic measures collected in HCP and ABCD was based on results from Chang et al. (Chang et al., 2018) where significant differences involving attention networks (DAN, VAN) and their connectivity with FPN and SMN networks were found to differentiate CWS from controls. Clinically, ADHD and subclinical attention deficits are commonly reported in stuttering (Donaher & Richels, 2012), but to date, there have been few studies investigating how neural networks supporting attention are affected in stuttering. Sustained attention in HCP is measured using the Short Penn Continuous Performance Test (SCPT) (Gur et al., 2010; Kurtz et al., 2001). Participants see vertical and horizontal red lines flash on the computer screen. In one block, they must press the spacebar when the lines form a number, and in the other block, they push the spacebar when the lines form a letter. The lines are displayed for 300 ms, followed by a 700 ms ITI. Each block contains 90 stimuli and lasts for 1.5 minutes. The equivalent sustained attention variable from the ABCD study is from the stop signal fMRI task (SST), corresponding to the total number of correct go trials across the entire task. The SST requires participants to withhold or interrupt a motor response to a “Go” stimulus when followed unpredictably by a signal to stop. Each of the two runs contains 180 trials. A further detailed description of this task can be found in Casey et al. (Casey et al., 2018).

Finally, to test whether big data improves classification performance, we compared the big data models to within-sample feature discovery models. Using big datasets from HCP and ABCD may be hurting our predictive performance due to differing age ranges of the populations they are drawn from and that there are unlikely any clinically diagnosed stuttering participants in these datasets. Instead of using the HCP or ABCD basis set, we embedded the dimensionality step within the stuttering dataset cross validation, and this model is called “In-sample top k=10”. We used the top k =10 components from the stuttering training set as predictors and compared them to the top k=10 HCP model and k=10 ABCD model. We did not repeat the pheno-corr method in-sample due to not having enough training set subjects to yield reliable correlations (Poldrack et al., 2020; Varoquaux, 2018; Varoquaux et al., 2017).

### Model Evaluation

The performance of our model was determined using the area under the curve (AUC) (Fawcett, 2006; Hand & Till, 2001). The “curve” reference in AUC corresponds to the receiver operating characteristic (ROC) curve, which plots the true positive rate versus the false positive rate. AUC is calculated on the test set within 10-fold cross validation, and the average AUC across folds is reported in Table 2 and Figure 2. Assessing the overall model statistical significance in our primary analysis framework is challenging due to the cross-validation procedure (10 different test-sets). While we do not want to rely on statistical significance for interpreting results, we recognize that overall model significance is helpful for determining that the prediction is meaningful (above chance accuracy). Following model evaluation and reporting methods in previous work (Sripada, Angstadt, Rutherford, Taxali, Greathouse, et al., 2020), a logistic regression model was fit within the whole sample to determine the overall statistical significance of each proposed feature selection method. The statistical significance of each full models’ predictors is shown in supplemental tables, and the overall model statistical significance is reported in the results section.

**Table 2.** **HCP & ABCD pheno-corr model performance** using variables from the NIH Toolbox and a sustained attention task. The number of significantly correlated components with phenotypes (p<0.01) are reported for each dataset along with the average AUC across a 10-fold cross validation framework.

| Phenotype | Subdomain | Ability | # of HCP components ($p < 0.01$) | HCP AUC | # of ABCD components ($p < 0.01$) | ABCD AUC |
| --- | --- | --- | --- | --- | --- | --- |
| Dimensional Change Card Sort | Executive Function | Cognitive Flexibility | 8 | 0.52 | 14 | 0.58 |
| List Sorting | Working Memory | Working Memory | 9 | 0.57 | 21 | 0.53 |
| Picture Vocabulary | Language | Vocabulary Knowledge | 13 | 0.54 | 17 | 0.59 |
| Pattern Comparison | Processing Speed | Processing Speed | 13 | 0.52 | 15 | 0.65 |
| Picture Sequence | Episodic Memory | Episodic Memory | 15 | 0.54 | 22 | 0.50 |
| Reading Recognition | Language | Oral Reading Skill | 18 | 0.52 | 18 | 0.58 |
| Fluid Intelligence<sup>2</sup> | - | - | 13 | **0.66** | 17 | **0.66** |
| Sustained Attention | - | - | 13 | 0.58 | 17 | 0.56 |

**Figure 2.**
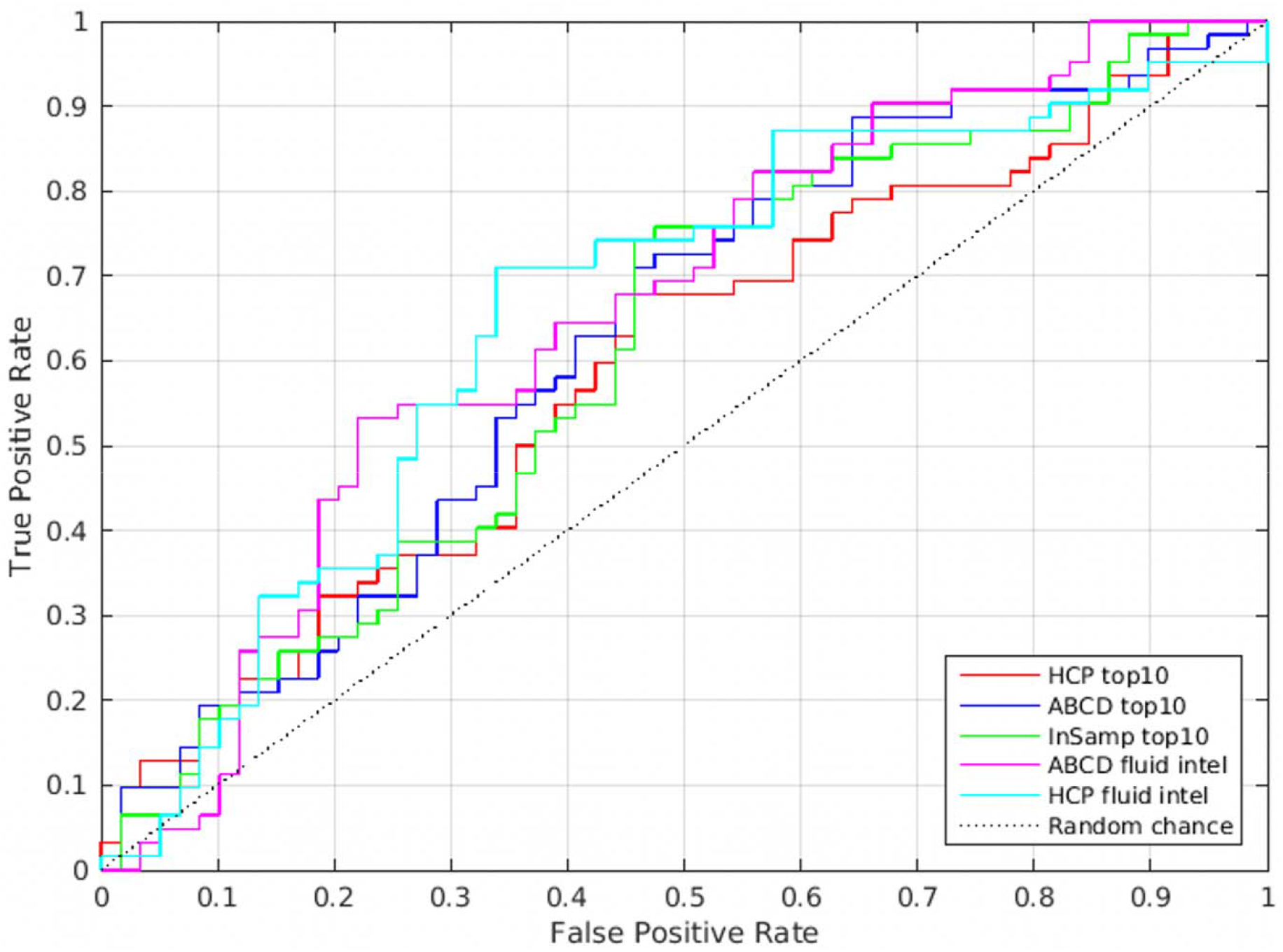
Whole Brain Receiving Operator Characteristic Curve plotting the false positive rate versus true positive rate for the top k=10 HCP, ABCD, and In-sample models and the fluid intelligence HCP and ABCD models. All of these models were tested using 10-fold cross validation logistic regression models. The diagonal dotted line shows chance accuracy. For most all subjects, the big data models (blue, red, cyan, magenta lines) are above the in-sample model (green line) meaning that big data models help improve the classification for most subjects in the clinical dataset.

### Visualization of Whole Brain Predictive Models

To help convey overall patterns across all components in a given BBS predictive model, we constructed “consensus” component maps, shown in Figure 3. We first fit a BBS model to the entire dataset consisting of all stuttering participants. We then multiplied each component map with its associated beta weight from this fitted BBS model. Next, we summed across all top k=10 components or pheno-corr components, yielding a single map for each model. The resulting map indicates the extent to which each connection is positively (red) or negatively (blue) related to the outcome variable of interest, stuttering status.

**Figure 3.**
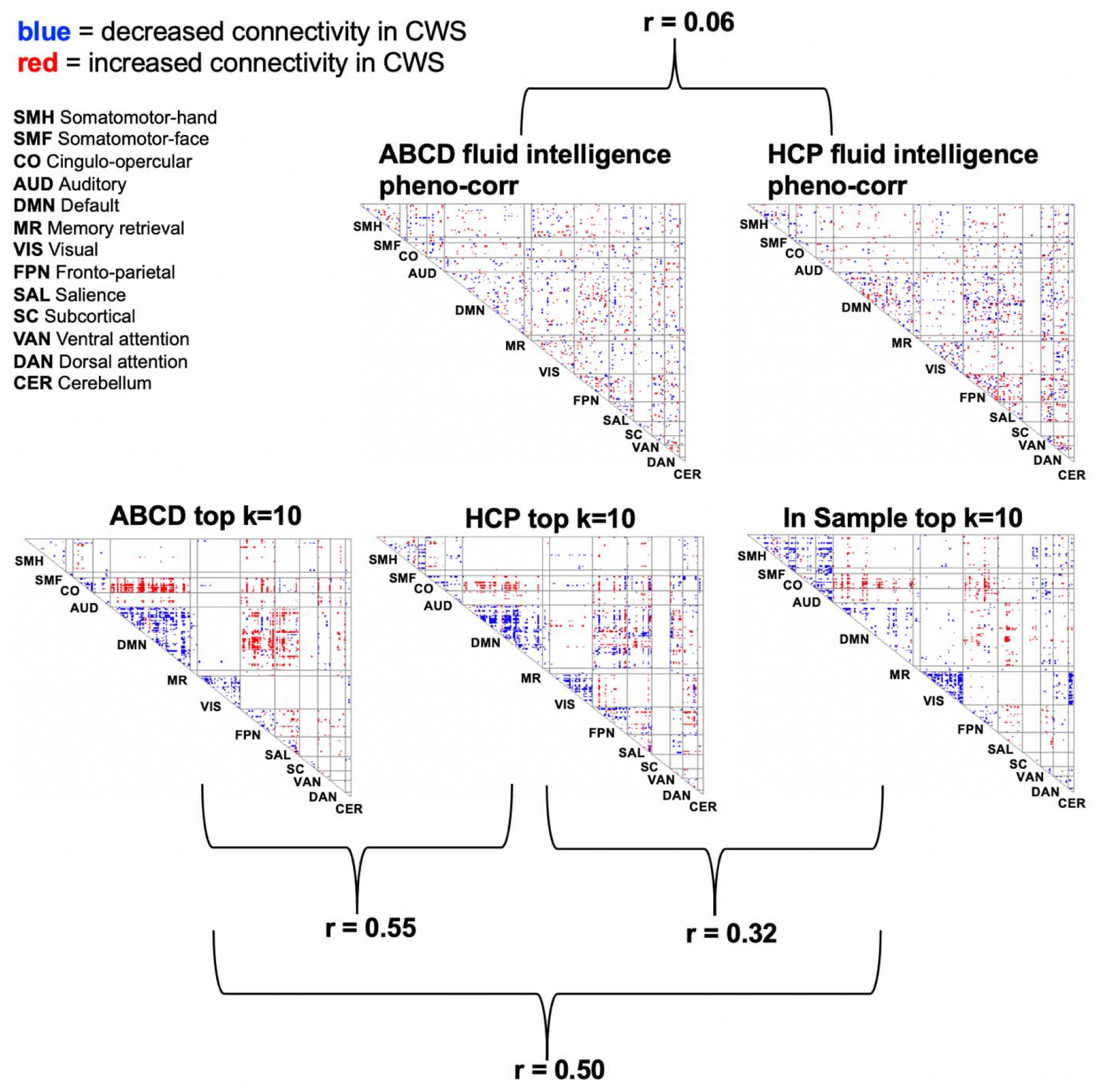
Consensus maps of connections that are predictive of stuttering status. The top k=10 models are highly similar across all three datasets (bottom row), and the pheno-corr models (top row) show unique, distributed patterns of connections that predict stuttering status. The correlation (r) between each of the maps is shown to quantify the overlap between each model’s predictive features.

### Interpretation Analysis: Keep Two Networks

To aid in the predictive model’s interpretability, we repeated all analyses using every network pair (one at a time) from the Power parcellation (Power et al., 2011). There are 13 brain networks in the Power parcellation, which results in 78 network pairs (13 choose 2). This allows us to observe which nodes, edges, and networks contribute most to the predictive model’s accuracy. For every network pair, for example, the FPN-DMN, all other nodes, and edges belonging to other networks are removed to create a much smaller connectivity matrix (that includes within-DMN, within-FPN, and DMN-FPN connections) for each subject and every step of the analysis (PCA matrix decomposition, component selection, and cross-validated logistic regression model fit) are performed on only the nodes and edges belonging to the FPN & DMN networks. This was repeated for all other network pairs. Figure 3A provides a visual intuition for the process of creating network-pair connectomes from the whole brain connectome. We used the top k=10 components HCP, ABCD, and In-sample models during the component selection step. For the pheno-corr models, we only used the HCP fluid intelligence model and ABCD fluid intelligence model due to their high prediction accuracy in the whole-brain results. Also, given that there are 78 network-pairs, and therefore 78 models to run *per phenotype,* we needed to limit our selection to produce succinct results.

### Data sharing

All code and data (that allows for it) is made available on GitHub (https://www.github.com/saigerutherford/bigdata-stuttering). The ABCD data does not allow raw or derivative data re-sharing and requires users to complete their own data use agreement on https://www.nda.nih.gov. A study has been created on NDA (DOI:10.15154/1520500) to track the included ABCD subjects in these analyses.

## Results

### Primary Analysis: Whole-Brain Connectome

#### Fluid intelligence pheno-corr model in ABCD and HCP achieved highest accuracy

The brain basis set predictive model successfully differentiated between children who stutter and healthy controls using resting-state connectivity patterns from out-of-sample healthy child (ABCD study) and adult (HCP study) datasets. The best performing whole-brain model used components related to fluid intelligence in the child (AUC_10fold-cv_ = 0.66; p-value_full-sample_ = 5.79e-5), and adult (AUC_10fold-cv_ = 0.66; p-value_full-sample_ = 2.25e-3) samples. Accuracy of other phenotype models from the NIH-Toolbox and sustained attention ranged from 0.5 (Picture Sequence – episodic memory) to 0.65 (Pattern Completion – processing speed). Table 2 summarizes the accuracy of all ABCD and HCP pheno-corr models, along with the number of features in each model (components significantly correlated with each phenotype), and the subdomains/abilities each phenotype represents. The supplemental tables contain the feature level statistical significance of all models.

### Top k=10 models suggest big data basis sets may improve accuracy

When comparing the HCP, ABCD and in-sample top k=10 basis sets, the ABCD (AUC_10fold-cv_ = 0.63; p-value_full-sample_ = 2.26e-3) and HCP (AUC_10fold-cv_ = 0.59; p-value_full-sample_ = 4.58e-3) models had slightly higher accuracy than the in-sample model (AUC_10fold-cv_ = 0.57; p-value_full-sample_ = 7.85e-3), suggesting that using big data to discover a brain basis set improves prediction performance, or at the very least does not decrease performance compared to in-sample feature selection. Interestingly, the top k=10 predictive models from separate datasets appear to leverage very similar brain connections. The correlation between the consensus maps of each model are strongly positive and are statistically significant. The HCP & ABCD consensus maps are correlated r=0.55, HCP & In-sample correlation of r=0.32, and ABCD & In-sample are correlated r=0.50. The pheno-corr fluid intelligence model showed less overlap between the HCP & ABCD models (r=0.06). The ROC curve for all dataset’s top k=10 and ABCD & HCP fluid intelligence pheno-corr whole brain models are shown in Figure 2 and the consensus map models for all dataset’s top k=10, and HCP & ABCD fluid intelligence pheno-corr models are shown in Figure 3.

### Interpretation Analysis: Keep Two Networks

#### Subset network basis sets reveal a fine-scale resolution of clinical data not detected in whole-brain models

When further interrogating which brain nodes, connections, and networks contributed the most useful information to our predictive model, we learned that the auditory and ventral attention networks, from the in-sample top k=10 model, contributed the most and yielded the highest accuracy (AUC_10fold-cv_ = 0.72). The salience and subcortical network pair from the in-sample top k=10 model was a close second (AUC_10fold-cv_ = 0.71). These results show that when using a reduced basis set (from just two brain networks, e.g., auditory & salience), the in-sample basis set outperforms ABCD & HCP. The results of all network pairs are visualized in Figure 4. This suggests that clinical variation has a fine-scale resolution that may be overlooked when searching the full connectome.

**Figure 4.**
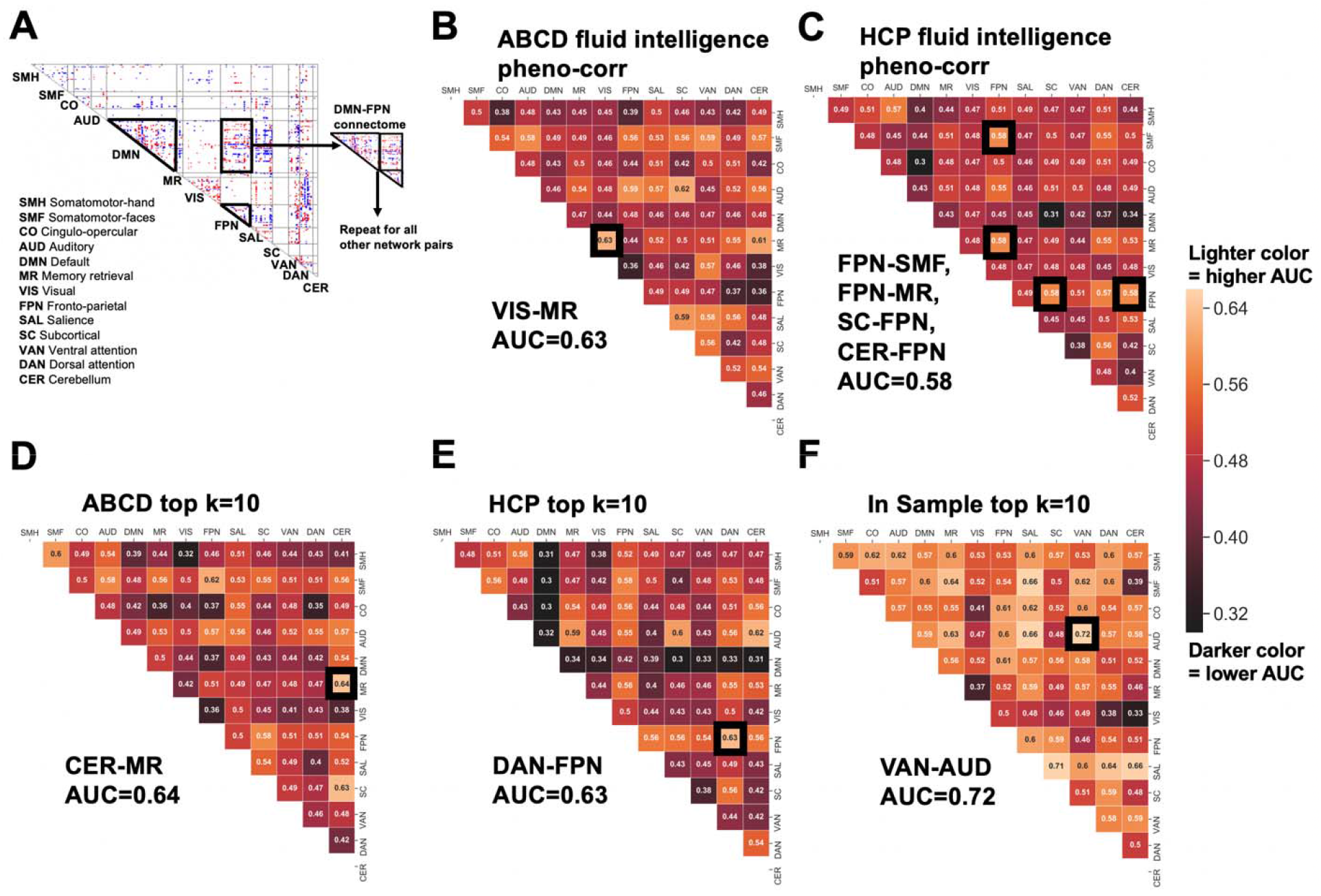
Keep Two Network Analysis. **A)** Visual intuition for how the network-pair connectomes (ex. DMN-FPN) are created. This process was repeated for all 78 network pairs. In panels B-F, a black box is placed around the most accurate (i.e., best-performing model) network-pair, and this network pair is also shown below the panel in larger, bolded text. The number represented in each box is the average AUC across 10-fold CV for that given network-pair. Lighter colors correspond to higher accuracy. **B)** ABCD fluid intelligence pheno-corr **C)** HCP fluid intelligence pheno-corr **D)** ABCD top k=10 **E)** HCP top k=10 **F)** In-sample top k=10

## Discussion

In this work, we determined whether a brain basis set developed in big data sets, HCP and ABCD, could be used to classify stuttering children from fluent peers. The results across several predictive models in this study suggest that big data can be transferred to smaller clinical datasets to help prediction performance. While the big data-based models provide a rather modest improvement in classification accuracy, it is important to remember that predicting behavior from biological data is a highly complex problem to solve, and accuracy is not the only important facet of predictive modeling. There are other benefits of leveraging big data such as testing true out of sample model fit to determine generalizability and exploration of brain components related to phenotypes that were not collected in smaller clinical samples.

Past neuroimaging research has pointed to a wide range of structural and functional deficits in speakers who stutter, encompassing aberrant auditory-motor cortical connectivity and basal ganglia-thalamocortical connections. The wide range of locations and connectivity patterns that differ in stuttering speakers may not be surprising, given that multiple neural systems’ deficits can have detrimental effects on fluent speech production. While the main behavioral manifestations of stuttering involve speech disfluency, many children who stutter also exhibit comorbid symptoms comprising subtle language, attention, and/or cognitive deficits. This finding is similar to that observed in other neurodevelopmental disorders where diagnostic categories overlap and are highly heterogeneous (Siugzdaite et al., 2020). Reflecting these views, there is a growing consensus for rejecting a “core-deficit hypothesis” in developmental disorders in favor of embracing the view that neurodevelopmental conditions can arise from complex patterns of relative strengths and weaknesses that may encompass multiple aspects of cognition and behavior (Astle & Fletcher-Watson, 2020). Therefore, we tested whether cognitive functions that are more reliably captured with big data (using methods such as brain basis set [BBS] modeling to derive basic “features” inherent in resting-state fMRI) could improve classification performance in a pediatric stuttering dataset. We expected that this approach would allow us to examine how the stuttering group differs in these basic features in complex ways in the context of whole-brain connectivity measures.

The whole-brain connectome results showed that the BBS models of fluid intelligence phenotype-correlation derived from the HCP and ABCD datasets achieved the highest accuracy in classifying stuttering children. The fluid intelligence measure is a composite score that encompasses scores from other tests administered as part of the NIH toolbox®, including Pattern comparison task (processing speed; information processing), list sorting working memory task (working memory; Categorization; Information Processing), picture sequence memory task (Visuospatial sequencing & memory), flanker task (Cognitive Control/Attention, and the dimensional change card sort task (Flexible thinking; concept formation; set-shifting) (Gershon et al., 2010; Luciana et al., 2018). Interestingly, phenotypes that were more directly related to language function, such as reading and picture vocabulary scores, did not perform as well in the pheno-corr models compared to the fluid intelligence and processing speed cognitive phenotypes. These results suggest the need to examine further inherent changes in these cognitive dimensions (processing speed, attention, working memory) for stuttering children and how they might interact with stuttering status.

Apart from the pheno-corr models, the top k=10 models from ABCD and HCP performed slightly better than the model based on the in-sample dataset. The consensus component maps, generated to convey overall patterns across all components in a given BBS predictive model, further showed the connectivity pairs in each model that contributed to predicting stuttering status. The top k=10 predictive models from HCP, ABCD, and the patient samples, appeared to leverage very similar brain connections. The correlation between each model’s consensus maps was strongly positive and statistically significant, especially between ABCD and the patient sample. The higher correlation found between consensus models derived from ABCD and the stuttering sample could be attributed to the more similar age distributions of these two samples as opposed to the HCP study, which included mostly adults. Network-level alterations predicting stuttering based on consensus models that were common across all three datasets included: within-network connectivity decreases in the default mode network (DMN), frontoparietal network (FPN), and visual network. Also, increased connectivity between DMN-cingulo-opercular (CO) networks and the FPN-CO networks predicted stuttering status. The DMN is hypothesized to implement emotion regulation and self-inspection; decreased DMN function may negatively affect adaptive emotion regulation (Schilbach et al., 2012). Decreased DMN function, when paired with decreased FPN function (affecting cognitive control) and overactive functioning of CO and ventral attention networks (VAN), have been implicated in anxiety disorders (Sylvester et al., 2012). The fact that stuttering was associated with decreased intra-network connectivity of both the DMN and FPN networks and at the same time increased connectivity of those same networks with the cingulo-opercular network is interesting given the CO network’s relevance to error sensitivity, detecting negative affect and pain (Shackman et al., 2011). Altered within-network functional connectivity in the CO network has also been linked to patients with social anxiety disorder (Liao et al., 2010) and tonic alertness, i.e., sustained attention (Sadaghiani & D’Esposito, 2015). The heightened connectivity between the cingulo-opercular network to both DMN and FPN networks in stuttering speakers might reflect the need for greater involvement of these two networks to achieve emotional regulation and cognitive control in the presence of error detection and conflict.

While social anxiety is commonly reported in adults who stutter, the same has not been consistently reported in children who stutter. Direct examination of emotional processing in children who stutter has been rare, though some studies have reported subtle differences in CWS in terms of autonomic nervous system responses to challenging speech tasks (nonword repetition; (Tumanova & Backes, 2019)) or emotionally stressful stimuli ((Jones et al., 2014; Walsh et al., 2019). The current BBS results suggest that brain networks linked to emotional regulation and their connectivity with cognitive control networks may differentiate children who stutter. More research is warranted in this area, especially related to understanding how these network connectivity patterns change due to persistence and stuttering recovery.

Altered network findings predicting stuttering status based on just the in-sample consensus model included: decreased within-network connectivity in the auditory and cerebellar networks and decreased connectivity between the auditory-somatomotor networks (SMF, SMH) and auditory- ventral attention networks (VAN). The in-sample model pointing to the auditory network and its connectivity with somatomotor networks is largely consistent with past literature in stuttering, where most studies have identified structural and functional deficits in auditory-motor integration. Decreased within-network connectivity in the auditory network may suggest a less ideal functioning of this network, representing a critical to interface with the speech motor region, necessary for developing and maintaining fluent speech control (Bohland & Guenther, 2006). Deficient auditory cortex function has been reported as one of the neural “signatures” of stuttering based on meta-analyses of neuroimaging stuttering literature (Brown et al., Budde et al.). The current results implicating decreased auditory-ventral attention network connectivity in children who stutter is relevant to findings reported in a previous study of childhood stuttering that also showed aberrant connectivity involving the attention networks, including the ventral attention network (Chang et al., 2018). The VAN includes parts of the ventrolateral prefrontal cortex and the temporoparietal junction and supports bottom-up attention, i.e., directing attention to newly appearing stimuli. The decreased connectivity between the VAN and auditory networks implicated in the in-sample consensus model for stuttering may suggest discordant function between these two networks. That is, an abnormally increased function of the ventral attention network may be linked to a tendency to direct attention to stimuli that suddenly appear rather than towards (auditory) stimuli that are currently the focus of the task to be performed, e.g., speech control. A prolonged influence of such aberrant auditory-VAN connectivity may cascade downstream to influence how the auditory network interfaces with the somatomotor networks, which could lead to inefficient integration of auditory-motor networks that are critical for supporting fluent speech control.

Another altered network finding that was specific to the in-sample consensus model was decreased within-cerebellar network connectivity. Given the cerebellum’s role in motor learning, error correction, movement timing, and past findings of aberrant function (De Nil et al., 2003) and structure (Connally et al., 2014; Sitek et al., 2016) in stuttering speakers, a further detailed examination of this structure in relation to stuttering is warranted. Apart from its motor-related functions, the cerebellum has connections with most parts of the cerebral cortex (Buckner, 2013) including the auditory cortex. It also connects to the basal ganglia and thalamus and has afferent connections from the olivary nucleus. The latter may have a role in detecting and processing somatosensory and auditory errors (Schweighofer et al., 2013). It would be of particular interest to examine how cerebellar network connectivity, both within-network and between network connectivity - change during development in normally developing children compared to children who stutter. Such investigations have the potential to reveal how the cerebellar function may modulate previously reported network alterations in stuttering and provide clues to how this may influence developmental changes that are linked to later persistence and recovery.

The keep- two network models were used to interrogate further which brain nodes, connections, and networks contributed the most useful information to our predictive model using a reduced basis set (from just two brain networks). Here, the in-sample basis set outperformed the models from both the ABCD and HCP data sets. The auditory and ventral attention network pairs appeared to contribute the most to the model and yielded an accuracy of (72%). The salience and subcortical network pair from the in-sample top k=10 model was the next highest contributor achieving 71% classification accuracy. Here, the auditory-VAN network connectivity is highlighted again, providing further confirmation of these networks’ importance in predicting stuttering status. The salience network is often equated with both the CO and VAN networks, with overlapping or adjacent structures comprising each of these networks. The salience network’s key structures include the anterior insula and the dorsal anterior cingulate cortex, although the striatal-thalamic loop is also functionally connected to the key structures and hence the salience network (Peters et al., 2016) The salience network’s function has been reported to include detecting and integrating sensory and emotional stimuli (V. Menon, 2015; Vinod Menon & Uddin, 2010), and attentional shifting that helps mediate the switch between internally-directed versus externally-directed attention (Uddin, 2015). In the consensus models discussed above, connectivity involving the salience network was altered for stuttering children, based on both ABCD and in-sample models. Networks showing altered connections with the salience network included the DMN and FPN, two networks linked to internally directed and externally directed cognitive control function, respectively. These results suggest that the efficiency of switching between internal and externally oriented tasks might be affected in children who stutter. Specifically, these inefficiencies may be reflected in suboptimal performance on externally oriented tasks such as speech production because internally oriented processing such as self-inspection is not fully switched off. A similar perspective is taken by the default network interference model (Sonuga-Barke & Castellanos, 2007), where diminished segregation between the DMN and other networks might allow the intrusion of DMN activity that causes inefficient functioning of task-positive processes (Zou et al., 2013) and lead to behavioral variability (Kelly et al., 2008; Poole et al., 2016). Altered connectivity of the salience network to the subcortical network in stuttering is also not surprising, given past findings pointing to aberrant thalamocortical loop function in stuttering physiology (Alm, 2004; Chang & Guenther, 2020; Craig-McQuaide et al., 2014).

## Conclusion

In sum, our findings show that using big data such as ABCD and HCP datasets to derive basic cognitive “features” provided superior models to classify children who stutter from age-matched controls. The results provide a significant expansion to previous understanding of the neural bases of stuttering that had previously been limited mainly to auditory and motor integration areas in the cortical and subcortical regions. In addition to auditory, somatomotor, and subcortical networks, the models built using big data highlight the importance of considering large scale brain networks supporting error sensitivity (cingulo-opercular), attention (ventral attention, salience), cognitive control (FPN), and emotion regulation/self-inspection (DMN) in the neural bases of stuttering. The results also suggest that while big data can identify whole-brain based connectivity alterations relevant to the disorder, these approaches might be best supplemented by detailed reduced-basis set modeling that further interrogates which brain nodes, connections, and networks contribute the most useful information to the predictive models. This latter approach indicated that the clinically specific stuttering dataset outperformed the big dataset derived models, possibly showing core disorder-specific networks that may be altered and are vulnerable for further modulation from other large-scale networks. This study is a first attempt to identify the brain basis features predictive of stuttering. The present findings offer insights into the neurophysiological basis of stuttering and pave the way for future studies that elucidate neural mechanisms for ultimately predicting the optimal treatment strategy and/or outcomes. The transfer learning framework introduced by this work builds an important connection between the clinical neuroscience and the big-data neuroscience communities.

## Abbreviations

ABCD: Adolescent Brain Cognitive Development study
ADHD: Attention Deficit Hyperactivity Disorder
AROMA: Automated Removal of Motion Artifact
AUC: Area under the Curve
BBS: Brain Basis Set
BOLD: Blood Oxygen Level Dependent
CO: Cingular-Opercular network
CER: Cerebellum network
CSF: Cerebral Spinal Fluid
CWS: Children Who Stutter
DAN: Dorsal Attention network
DMN: Default Mode network
EPI: Echo Planar Imaging
FIX: FIMRIB’s ICA-based Xnoiseifier
FD: Framewise Displacement
FSL: FIMRIB Software Library
fMRI: Functional Magnetic Resonance Imaging
HCP: Human Connectome Project
ICA: Independent Component Analysis
MNI: Montreal Neurological Institute
M R: Memory Retrieval network
PCA: Principal Component Analysis
ROC: Receiver Operator Characteristic
ROI: Region of Interest
rsfMRI: Resting-State Functional Magnetic Resonance Imaging
SCPT: Short Continuous Performance Task
SMN: Somatomotor network
SPM: Statistical Parametric Mapping
SSI: Stuttering Severity Instrument
VAN: Ventral Attention network
VIS: Visual network

## ACKNOWLEDGEMENTS

Data used in the preparation of this article were obtained from the Adolescent Brain Cognitive Development (ABCD) Study (https://abcdstudy.org), held in the NIMH Data Archive (NDA). This is a multisite, longitudinal study designed to recruit more than 10,000 children age 9-10 and follow them over 10 years into early adulthood. The ABCD Study is supported by the National Institutes of Health and additional federal partners under award numbers U01DA041022, U01DA041028, U01DA041048, U01DA041089, U01DA041106, U01DA041117, U01DA041120, U01DA041134, U01DA041148, U01DA041156, U01DA041174, U24DA041123, and U24DA041147. A full list of supporters is available at https://abcdstudy.org/nih-collaborators. ABCD consortium investigators designed and implemented the study and/or provided data but did not necessarily participate in analysis or writing of this report. This manuscript reflects the views of the authors and may not reflect the opinions or views of the NIH or ABCD consortium investigators. The ABCD data repository grows and changes over time. The ABCD data used in this report came from NDA Study 576, DOI 10.15154/1412097, which can be found at https://nda.nih.gov/abcd/query/abcd-interim-annual-release-1.1.html. HCP <u>S1200</u> data release was used in this study.

## Author Contributions

Conceptualization: SR, MA; Methodology: SR, MA; Formal Analysis: SR, MA; Data Curation: SR, MA; Writing – Original Draft: SR, SC; Writing – Reviewing and Editing; SR, MA, CS, SC; Visualization: SR; Supervision: SC, CS; Funding Acquisition: SC, CS.

## Footnotes

1 The Stuttering Severity Instrument (SSI-4) was used to examine frequency and duration of disfluencies occurring in the speech sample acquired from each child who stutters. The SSI composite score incorporates frequency and duration of stuttered speech, as well as any physical concomitants associated with stuttering (Riley & Bakker, 2009). To be classified as a child who stutters, they needed to score in the very mild or higher range on the SSI composite score. For borderline cases, parent’s expressed concern of stuttering and clinician (certified Speech-Language Pathologist) impression confirming stuttering status were considered in making the determination of stuttering status.

2 Fluid Intelligence is a Cognition Composite Score that includes DCCS, Flanker, Picture Sequence Memory, List Sorting, and Pattern Comparison measures.

